## Supplemental Figures (Revised) for "A modular platform to display multiple hemagglutinin subtypes on a single immunogen"

**SUPPLEMENTARY MATERIALS:**

Figs S1-S7

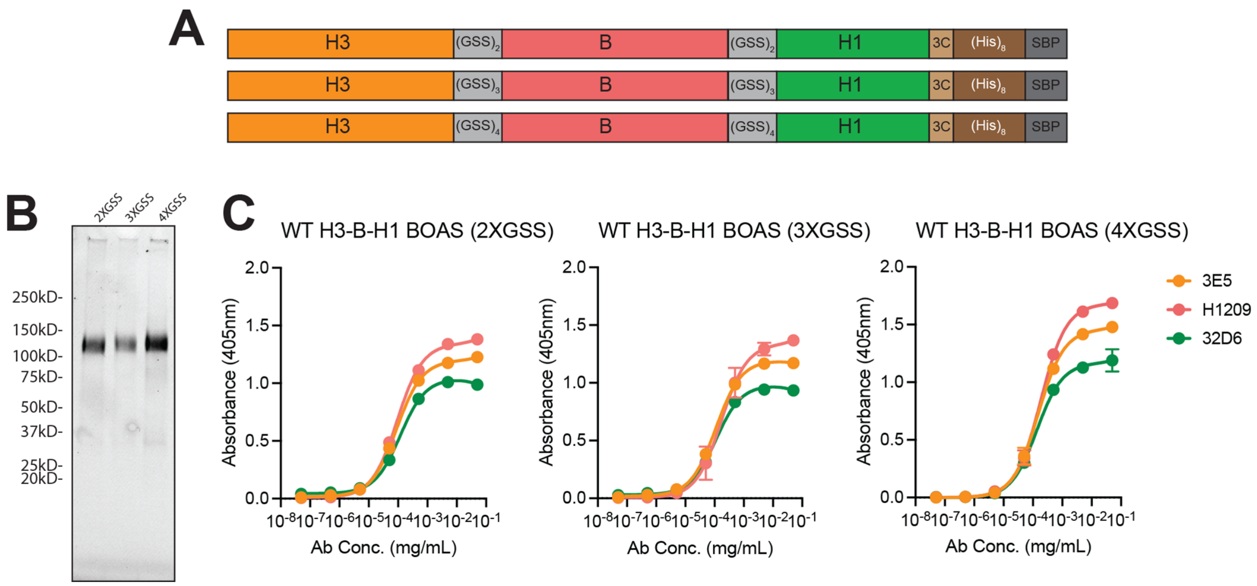


**Supplemental Figure 1: Adjusting flexible linker lengths on BOAS A)** Construct diagram of H3-B-H1 3mer BOAS with 2X, 3X, and 4X Gly-Ser-Ser linkers. **B)** SDS-PAGE of 3mer BOAS with 2X, 3X, and 4X GSS linkers from left to right. **C)** ELISA with subtype specific antibodies detecting H1 (green), H3 (orange), and B (salmon) subtypes on 2X, 3X, and 4X GSS linker BOAS.

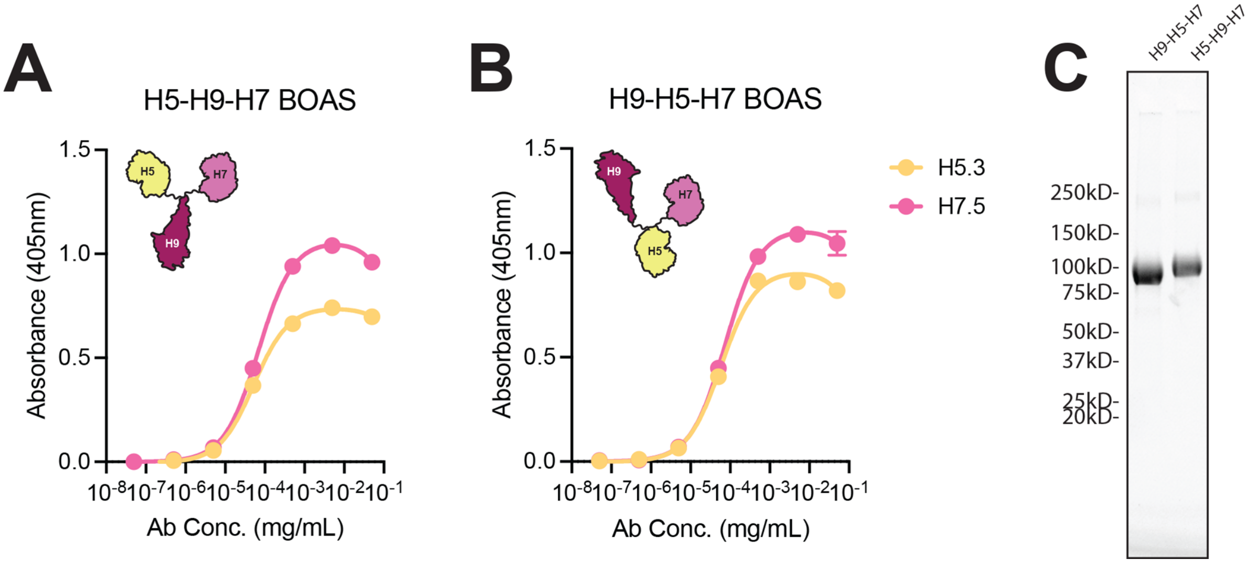


**Supplemental Figure 2: Swapping 3mer order on BOAS** Determining effect of switching order of BOAS heads on 3mer expression and ELISA. **A, B)** ELISA with H5 (yellow) and H7 (pink) subtype specific antibodies on BOAS with H5-H9-H7 (A) or H9-H5-H7 **B)** in order from N to C terminus. **C)** SDS-PAGE of H9-H5-H7 and H5-H9-H7 BOAS.

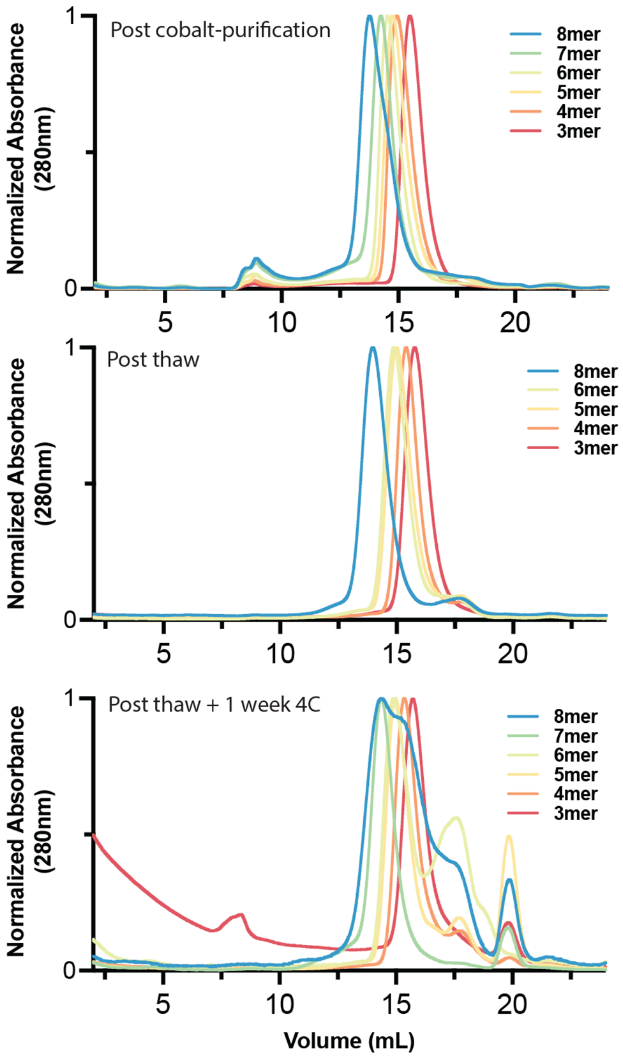


**Supplemental Figure 3: Size Exclusion Chromatography Traces of purified BOAS immunogens over time** – Traces of purified 3mer, 4mer, 5mer, 6mer, 7mer, and 8mer BOAS following cobalt purification (top), immediately thawed from -80C storage (middle), and post-thaw following 1 week of storage at 4C. Approximately 100µg of protein was injected onto a Superose 6 10/300 column in PBS, and each trace shows a normalized 280nm signal.

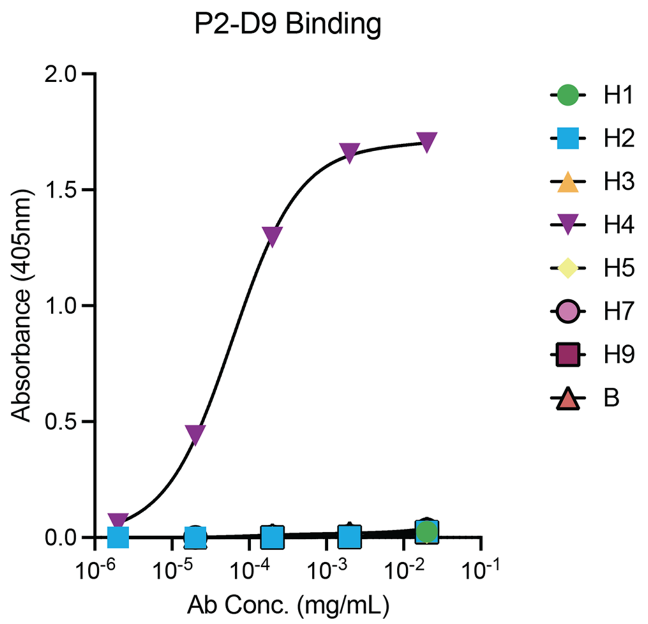


**Supplemental Figure 4: P2-D9 specificity for H4** ELISA assay to determine binding of P2-D9 to H1, H2, H3, H4, H5, H7, H9, and B influenza hemagglutinin (HA) trimers. P2-D9 only detectably engages H4/New Brunswick/2010 HA.

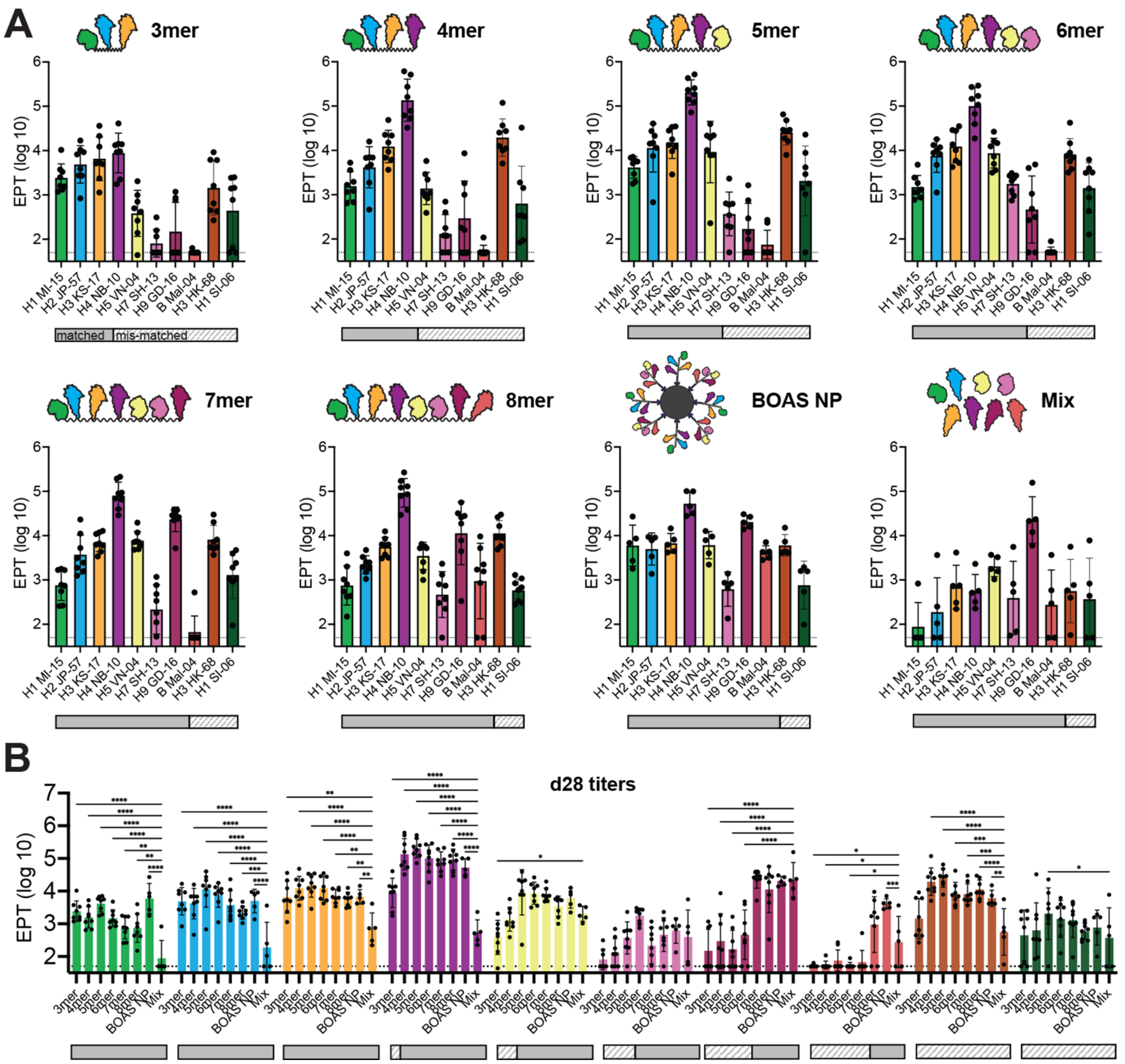


**Supplemental Figure 5: Day 28 Serum Reactivity from BOAS and BOAS NP cohorts-** A) Serum titers to full-length HA trimers elicited by 3mer-8mer BOAS and BOAS NP at d28. Solid bars below each plot indicate a matched sub-type, and striped bars indicate a mis-matched subtype (i.e. not present in the BOAS). Data points are serum EPTs from individual mice (n=8 for BOAS; n=5 for BOAS NP and Mix) and error bars are +/- 1 s.d. B) Serum reactivity to each matched and mis-matched individual full-length HA component of BOAS (H1/MI/15 light green, H2/JP/57 blue, H3/KS/17 light orange, H4/NB/10 purple, H5/VN/04 yellow, H7/SH/13 pink, H9/GD/16 maroon, B/Mal/04 salmon, H3/HK/68 dark orange, and H1/SI/06 dark green) by 3mer-8mer BOAS, Mix, and BOAS NP cohorts. Solid bars below each plot indicate a matched sub-type, and striped bars indicate a mis-matched subtype (i.e. not present in the BOAS or NP). * = $p<0.05$, ** = $p<0.01$, *** = $p<0.001$, **** = $p<0.0001$ as determined by Kruskal-Wallis test with multiple comparisons relative to the mix control group. * = $p<0.05$, ** = $p<0.01$, *** = $p<0.001$, **** = $p<0.0001$ as determined by a Kruskal-Wallis test with Dunn's multiple comparison post hoc test.

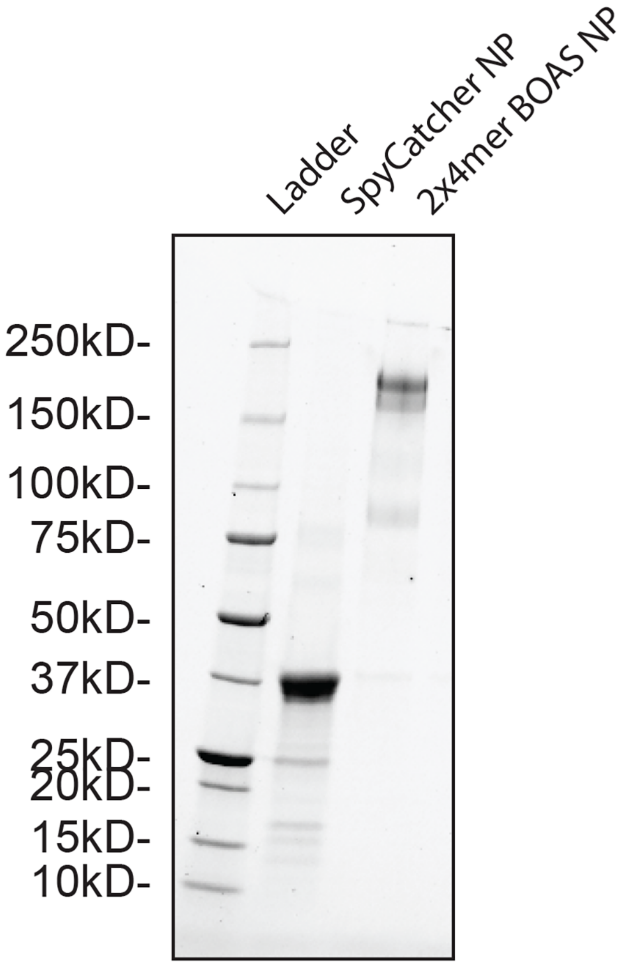


**Supplemental Figure 6: Nanoparticle conjugation efficiency SDS-PAGE –** SDS-PAGE gel of N-terminally fused SpyCatcher Ferritin (middle lane), and BOAS conjugated ferritin NP (right lane).

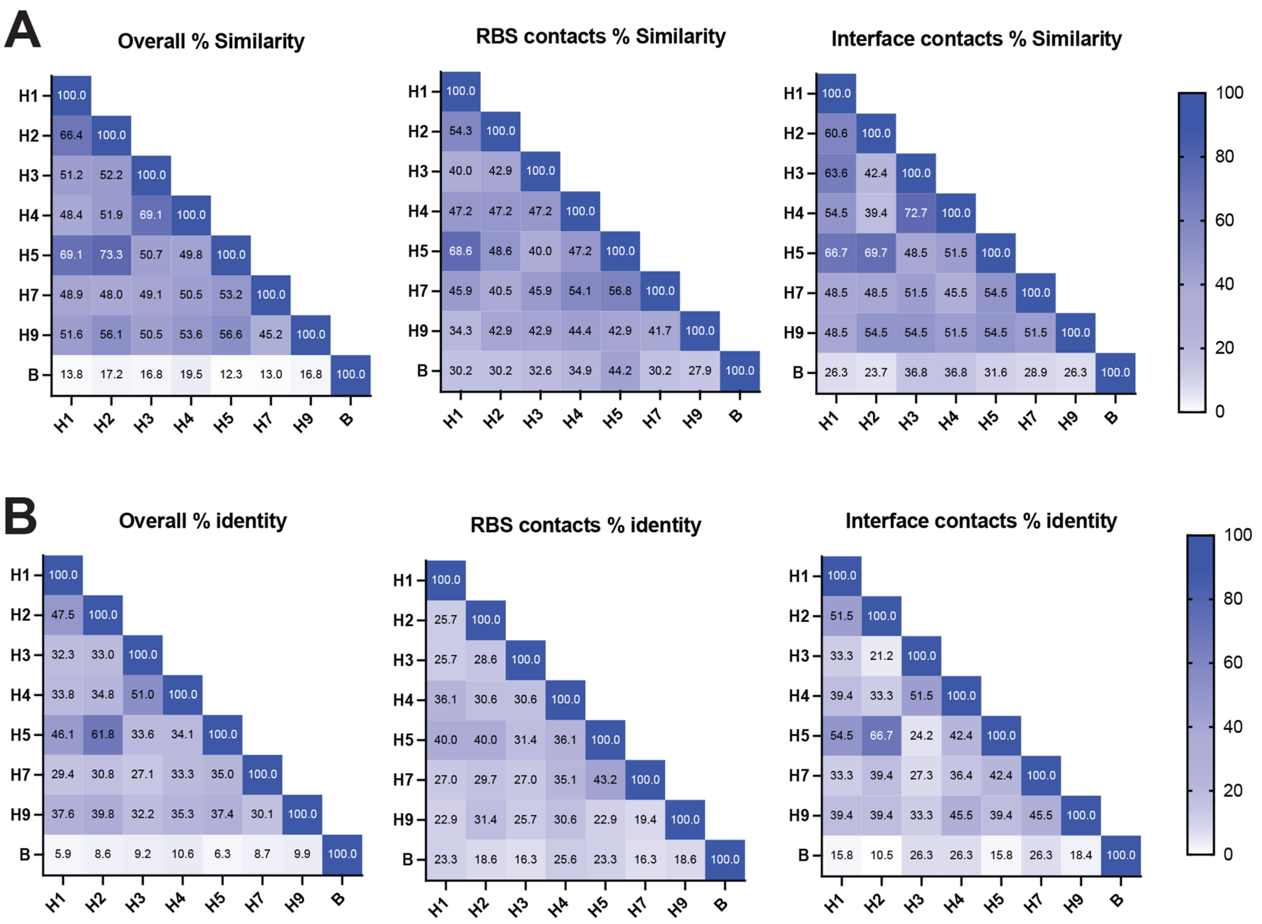


**Supplemental Figure 7: Sequence identity matrices of BOAS components** – Comparing sequence identity of eight BOAS components H1/Michigan/45/2015, H2/Japan/305/1957, H3/Kansas/14/2017, H4/New Brunswick/00464/2010, H5/Vientam/1203/2004, H7/Shanghai/01/2014, H9/Guangdong/MZ058/2016, B/Malaysia/2506/2004. Blosum45 **A)** similarity **B)** or amino acid identity matrices of total head domain amino acid sequence (left), receptor binding site (RBS) contacts (center), or interface contacts (right).
